# The pangenome of the human fungal pathogen *Cryptococcus neoformans* reveals that subtelomeres are hotspots for gene content variation

**DOI:** 10.64898/2026.10.27.762246

**Authors:** Claire E. Reichardt, R. Blake Billmyre

## Abstract

*Cryptococcus neoformans* is a human fungal pathogen that causes cryptococcal meningitis, primarily in individuals with advanced HIV disease. The *C. neoformans* species is composed of four unique molecular types: VNI, VNII, VNBI, and VNBII, with thousands of global isolates within these groups. While previous descriptions of these molecular types have focused on nucleotide divergence between shared genes, there has been limited examination of variation in gene content between isolates. Here, we define the genome content diversity within a collection of 383 *C. neoformans* isolates spanning all four molecular types. To do so, we constructed a pangenome through *de novo* assembly and annotation of short-read Illumina sequences. We defined 6,564 representative ortholog groups of the predicted proteins leveraging high-quality data from reference strain H99. We classified ortholog groups as either core (93.8%) or accessory (6.2%) based on their presence across the species. We next used long-read assemblies for a subset of our isolates to define the predicted accuracy of our classifications and account for errors in assembly or annotation of short-read sequences. We determined that while a subset of accessory genes are segregating across the entire species, many accessory genes are found only within specific molecular types, suggesting limited gene flow between groups. Additionally, we found many accessory genes are poorly conserved within the *C. neoformans* species but share high sequence similarity to genes found in related Cryptococcaceae species, suggesting horizontal gene transfer. We also found that accessory genes were enriched within subtelomeric regions, implicating the subtelomeres in gene content variation between isolates. We show examples of accessory genes located within subtelomeres of *C. neoformans* isolates that were conserved with subtelomeric proteins of other Cryptococcaceae species. Our pangenomic analysis of *C. neoformans* gives species-level insights into its biology, potentially illuminating genetic contributions to pathogenesis.

## INTRODUCTION

Invasive human fungal pathogens affect an estimated 6.5 million people each year[1]. One such pathogen is *Cryptococcus neoformans*, the causative agent of cryptococcal meningitis which causes 147,000 deaths annually[1]. Outside of patients, *C. neoformans* naturally occurs in the environment on trees or in pigeon guano[2]. From there, spores or desiccated yeast cells can be inhaled and cause invasive infections in immunocompromised patients. Cryptococcosis is an AIDS-defining illness accounting for 19% of deaths of AIDS patients[3]. While disease is most common in conjunction with the HIV/AIDS epidemic in equatorial and southern Africa, *C. neoformans* can be isolated around the world both from the environment and from patients.

Increased availability of whole genome sequencing has exponentially expanded the number of available sequences for a given species. The bloom in sequencing accessibility has driven a new type of genomics: pangenomics. A pangenome can be simply described as the collection of genes within a species, a replacement for singular reference representatives. The first pangenome described was in the bacterium *Streptococcus agalactiae* in 2005[4]. Tettelin et al. described two components to a pangenome: the core genome and the dispensable genome[4,5]. Core genes are conserved throughout the population while dispensable, or accessory, genes are present in some but not all members of the population. The core genome is thought to be important for essential biological functions[6] while the dispensable genome is important for survival in environmental niches[5]. Defining the core and dispensable genomes for a species can uncover population dynamics and environmental pressures driving genetic adaptation and niche specialization. Since the first described pangenome, pangenomics has rapidly expanded past prokaryotes to complex eukaryotic organisms including plants[7], humans[8], and fungi[6,9–14].

The first fungal pangenome was built using 1,011 isolates for the model yeast *Saccharomyces cerevisiae*[13]. The *S. cerevisiae* pangenome was defined to have 7,796 open reading frames within the population, of which 4,940 (63%) were core and 2,856 (37%) were accessory[13]. Along with genome-wide association studies (GWAS), the *S. cerevisiae* pangenome revealed the plurality of phenotypic variation was explained by copy number variation (36.8%), not single nucleotide polymorphisms (4.49%)[13]. Uniquely, the *S. cerevisiae* pangenome defined the subtelomeres as a hotspot for novel genes due to horizontal gene transfer from related species[13].

Since the *S. cerevisiae* pangenome, a variety of pangenomes have been built for fungal pathogens including *Aspergillus fumigatus*[9,11]. The *A. fumigatus* pangenome contains a much larger number of genes within its dispensable genome, with only 51.97% of ortholog groups classified as core[9]. One potential explanation for this difference is that the *A. fumigatus* genome contains large transposons, named *Starships*, that move approximately 16% of the accessory genome in *A. fumigatus*[15]. In comparison, the *Candida parapsilosis* pangenome was mostly conserved, with only 1.2% of the pangenome being accessory genes[10]. Despite its low gene diversity, large phenotypic variation is still present within the *C. parapsilosis* population[10]. No pangenome constructed for a pathogenic fungal species has reported similar subtelomeric accessory gene enrichment that was observed in the original *S. cerevisiae* pangenome.

*C. neoformans* isolates have been described as derived primarily from clonal expansion, with a large majority of isolates being *MAT*α[16,17]. *C. neoformans* isolates share an average nucleotide identity of 97.7% within the species. As a population, *C. neoformans* can be split into four molecular types: VNI, VNII, VNBI, and VNBII[16,17]. Molecular types VNI and VNII are thought to be globally distributed, while VNBs (VNBI and VNBII) have been previously restricted to Botswana[16]. VNB molecular types also have a higher proportion of *MAT***a** isolates than VNI or VNII, driving a lower level of clonality within the VNB subtypes[16]. Generally, VNB isolates are more genetically diverse than VNI isolates[16,18]. The *C. neoformans* reference strain, H99 (VNI), was originally isolated from the cerebral spinal fluid (CSF) of an infected patient[19]. H99 is highly virulent in rabbits and mice, making it an ideal strain for studying cryptococcal meningitis[19]. Since its original isolation, H99 (H99O) has gone through a series of genetic changes and passages to make several independent lineages[20]. One lineage in particular, H99S, was passaged through rabbit models to generate a strain with increased virulence[20]. H99 also has a unique chromosomal composition compared to other wild isolates, where chromosomes 3 and 11 have undergone a large reciprocal translocation[21]. While not thought to change virulence, this chromosomal rearrangement affects carbon metabolism and signal transduction as two genes that were located at the breakpoints were disrupted (*YPH2* and *TGR1*)[21]. H99 and its derivative strains, including KN99[22], have been valuable reference strains for the field but do not adequately represent the diversity present in the *C. neoformans* population.

Independent isolates of *C. neoformans* often exhibit a diverse array of phenotypes. Different molecular types have vastly different phenotypic traits including variation in melanization capability and oxidative stress response[17]. However, neither antifungal drug responses nor patient outcome are linked to molecular type or geographic location[17,23]. Molecular types can be further defined by their sequence types (ST) using multilocus sequence typing (MLST). Seven loci have been defined for MLST analysis to separate genetic background in *C. neoformans* including *CAP59*, *GPD1*, *LAC1*, *PLB1*, *SOD1*, *URA5* and the *IGS1* regions[24]. ST93, which is part of VNI, for example, is prominent in clinical cases in Uganda (60% of isolates)[23,25], Brazil (85% of isolates)[26,27], and India (71% of isolates)[28,29]. GWAS has identified SNPs in ST93 isolates that contribute to isolate-specific virulence in mice[30,31]. One of those genes was *ITR4*[30,31], an inositol transporter located on the subtelomere of chromosome 14. Taken together, we know that *C. neoformans* isolates differ dramatically in phenotype. However, previous studies have primarily looked for the source of this variation using a reference-strain-based framework, restricting their ability to link phenotype to causative differences in gene content. As a result, we have a limited understanding of not only when phenotypic differences are caused by gene content changes, but also how much variation in gene content exists in the species. Therefore, we sought to establish a broad pangenome capturing the complex genomic diversity of *C. neoformans* using an existing collection of sequenced isolates[17].

To do so, we *de novo* assembled and annotated 383 unique *C. neoformans* isolates to generate orthologous gene clusters. To control for errors introduced via use of short read assemblies or variation in gene annotation, we validated the presence of ortholog groups in long-read assemblies to generate accuracy scores for each gene cluster, resulting in an estimated accuracy of 99.9% for core ortholog groups. We found high conservation of genes throughout the population, with a total of 6,157 core and 407 accessory ortholog groups, a subset of which were found in multiple copies. Several accessory genes were restricted to a singular molecular type, with VNII being the most unique. However, despite previous reports of clonality, we found a number of accessory genes that were segregating in multiple molecular types. We found that accessory genes were enriched within subtelomeric regions of the genomes of long-read assemblies of isolates, similar to past observations in *S. cerevisiae*[13] but in contrast with other fungal pangenomes. We found a large portion of our accessory genes shared sequence similarity to related *Cryptococcus* and *Kwoniella* species. Overall, we concluded that *C. neoformans* carries most of its genomic content variation within the subtelomeres.

## RESULTS

### Limited accessory genes in the *Cryptococcus neoformans* pangenome

To capture the genomic content diversity of *Cryptococcus neoformans* as a species, we assembled and annotated 387 unique isolates previously Illumina sequenced as part of the *C. neoformans* Strain Diversity Collection from Desjardins et al.[17] (Figure 1, Table S1). Our assemblies contained an average of 108 contigs per genome and an average scaffolded N50 of 1,449,679 base pairs (Table S1). Our initial assemblies had an average BUSCO score of 97.32% (reference H99 BUSCO= 97.1%[32]) (Table S1). We filtered and removed four poorly assembled genomes based on low BUSCO scores or high numbers of contigs (BUSCO < 96% or contigs > 200). Our final genome set included 184 VNI, 15 VNII, 121 VNBI, and 63 VNBII isolates (total n=383). We broadly classified into two categories with 284 isolates from a clinical source (i.e. CSF, lung, sputum, blood) and 99 from environmental (i.e. pigeon guano, tree, soil) (Figure 2, Table S1).

**Figure 1.**
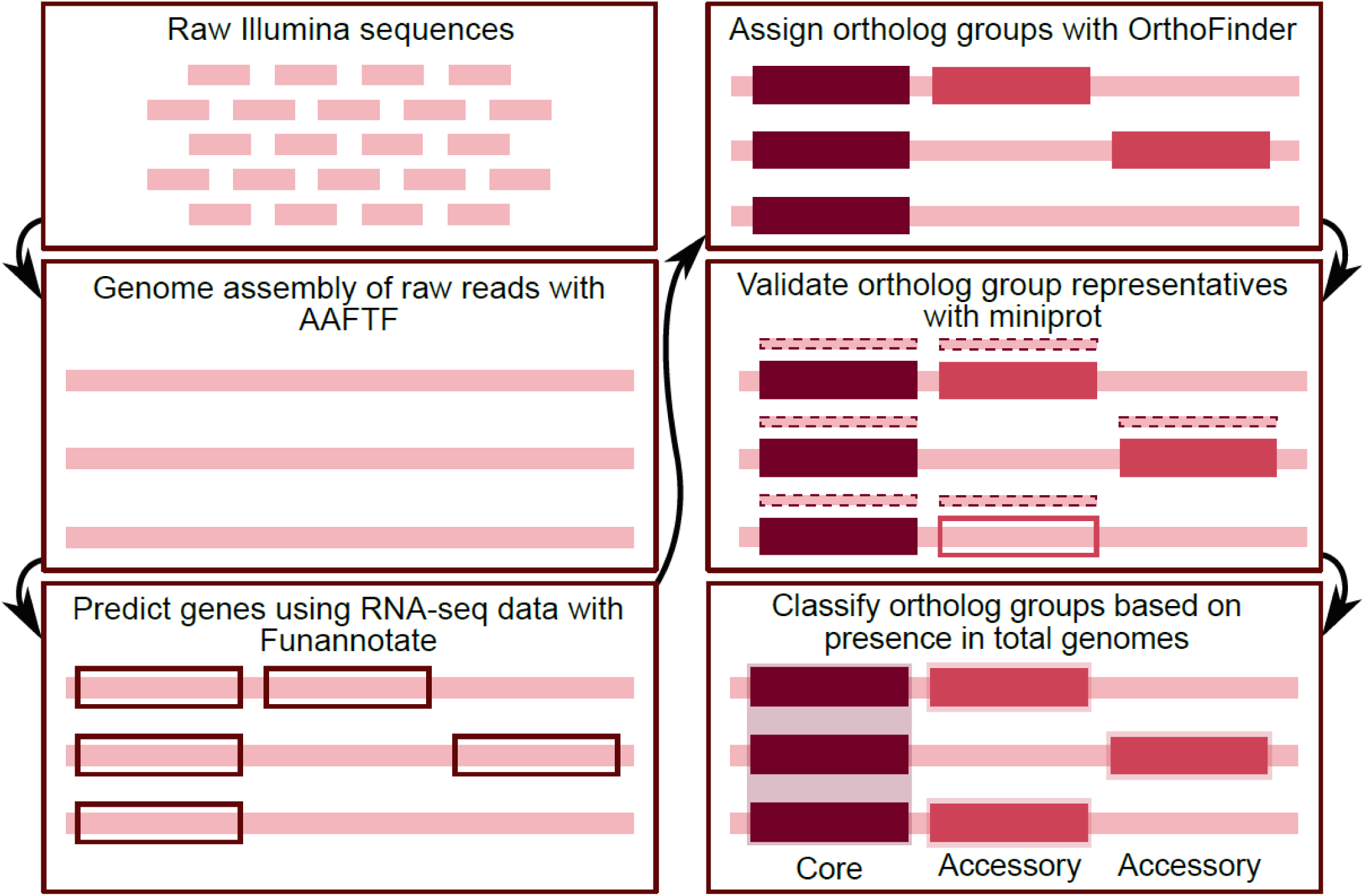
Pangenome assembly workflow. Publicly available raw Illumina reads from 387 diverse isolates were assembled using AAFTF and scaffolded using RagTag and a H99 reference genome. Scaffolds were annotated using Funannotate using RNA-seq data from reference H99. Predicted proteins were then fed into OrthoFinder to generate ortholog groups. One representative protein from each ortholog group was then mapped again to scaffolded assemblies to further annotate genomes. Using our final annotated proteins, we made two groups of ortholog groups: core (present in >95% of all genomes) and accessory (present in <=95% of all genomes).

**Figure 2.**
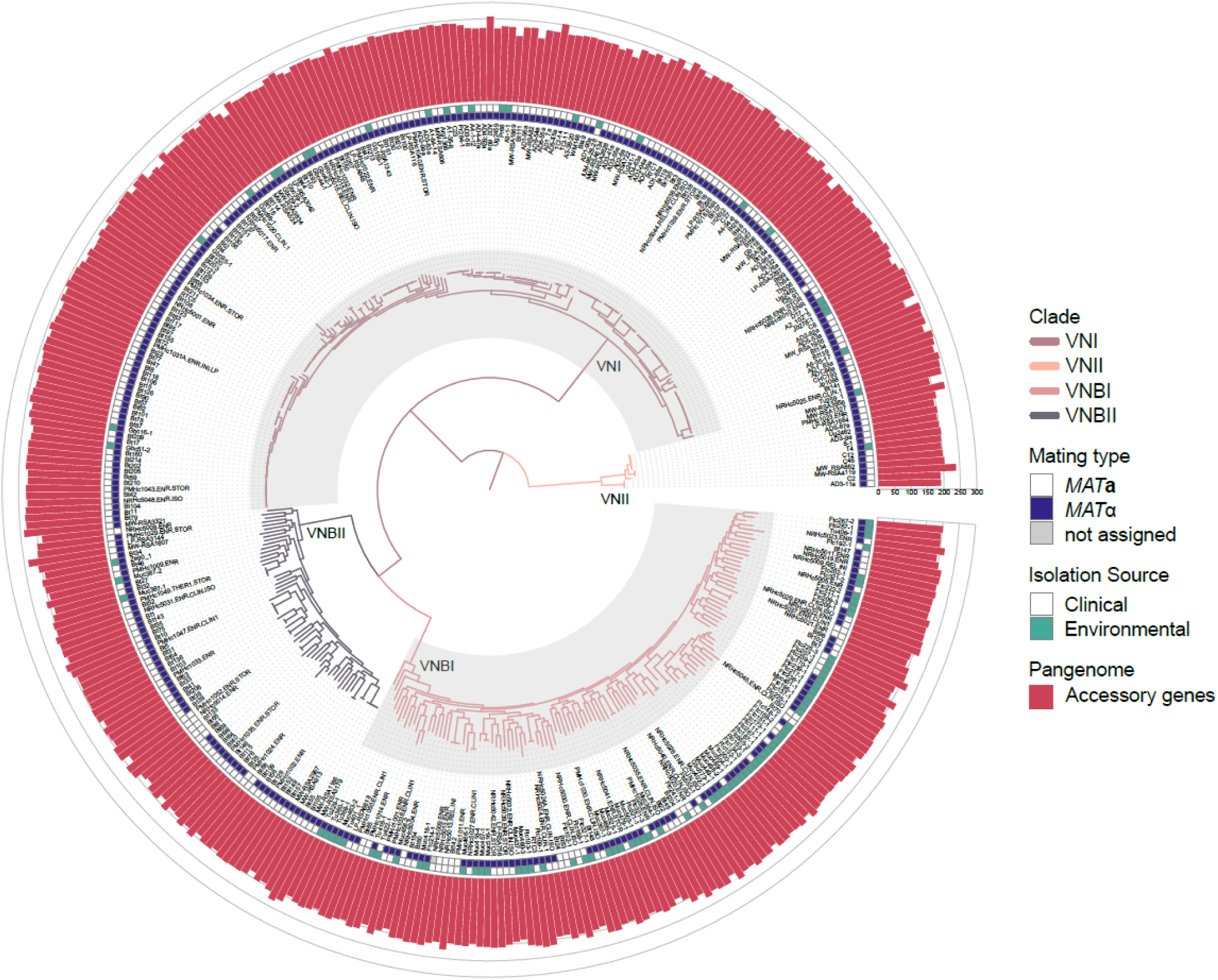
Accessory genes are widely distributed. Phylogenetic tree of 383 non-reference strains and their associated accessory gene counts. Tree was calculated by Desjardins et al. 2017 and rooted on VNII. First layer shows mating type as *MAT***a** (white) or *MAT*α (dark blue), and the second layer shows isolation source as clinical (white) or environmental (teal). Finally, pink bars show the total number of accessory genes called for each genome.

We then *de novo* predicted transcripts using Funannotate. This resulted in an average of 7,383 annotated proteins per genome. Our predicted proteins per genome was slightly higher than the annotation of reference strain H99 which was first annotated to have 6,967 proteins [20]. We preliminarily grouped proteins into ortholog groups. We followed *de novo* prediction by using representative sequences from each preliminary ortholog group to identify annotation gaps with miniprot. We defined annotation gaps as areas of assemblies that were highly similar to preliminary ortholog groups and had no annotation assigned yet. We then clustered identified genes using all by all comparisons with OrthoFinder supplemented with manual clustering and identified a total of 6,564 ortholog groups, including 6,157 core (93.8%) and 407 accessory ortholog groups (6.2%) (Figure 2, 3a). We assigned each ortholog group a sequential CN number (ie., CN00001-CN06564). Some CN groups contained multiple genes from a single isolate if they were multicopy (2,963 ortholog groups, Table S2). We identified only three singleton ortholog groups (ortholog groups that were present in only one isolate’s genome) in our pangenome. Therefore, we classified singletons as accessory genes instead of their own unique category.

To validate that accessory ortholog groups were truly missing from genomes rather than not assembled or annotated from short-read data, we benchmarked our results against long-read assemblies for a subset of the genomes in our dataset. To do so, we generated long-read sequences and *de novo* assembled 9 isolates and acquired another 25 existing assemblies from NCBI[33–39] (Table S3). Our long-read dataset included a total of 12 VNI, 3 VNII, 8 VNBI, and 11 VNBII isolates (Figure S1). These 34 assemblies had an average assembly size of 14.64 linear contigs (for 14 linear chromosomes) and an average contig N50 of 1,394,308 base pairs (not including mitochondrial genome) (Table S3). We used BUSCO to estimate the completeness of our assemblies and found an average score of 97.4%, comparable to that of our short-read assemblies. We annotated the long-read assemblies using miniprot to align our short-read assembly annotations to the long-read assemblies. We then calculated accuracy using long-read assemblies for a subset of the genomes in our dataset (Figure S2a). We predicted an average accuracy rate of 99.9% for core and 92.9% for accessory ortholog groups (Figure S2b). Core ortholog groups’ accuracy was more affected by false positives (called in short-read but not mapped in long-read assemblies, mean FPR=0.14) while accessory ortholog groups’ accuracy was primarily affected by false negatives (not called in short-read but mapped in long-read assemblies, mean FNR=0.13) (Figure S2b). Notably, ortholog groups that were present in fewer genomes were more likely to have a lower accuracy (Figure S2c).

Our comparison of long-read to short-read assemblies highlighted errors in the assembly process. For Muc525-1, we found that our short-read assembly generated a second tandem insertion of a 125 kb region near the left arm of chromosome 9, likely caused by repeats flanking the region. This region is inverted between *C. neoformans* and *C. deneoformans*, making it notably difficult to assemble with short reads[33]. As a result, approximately 44 proteins were annotated twice in Muc525-1 by Funannotate. Erroneous changes in gene copy numbers could be caused by instances like in Muc525-1, where short-read assemblies fail to properly capture the structure of repetitive regions.

A large proportion of the ortholog groups (n=5574, 84.9%) were called in all of the 384 genomes in our short-read assembly set (Figure 3b). We found 3,601 single copy ortholog groups present in every genome (Table S2). For accessory genes, we found that VNI genomes had a higher number of accessory genes called compared to other molecular types, resulting in a bimodal distribution for the entire species (Figure 3c). Upon comparing the assembly statistics and accessory gene numbers between molecular types, we found that VNI genomes had less fragmented assemblies, likely leading to a higher number of predicted accessory genes (Figure S3). This is most likely due to reference biases as we used the H99 (VNIb) genome and transcriptome to scaffold contigs and annotate proteins.

**Figure 3.**
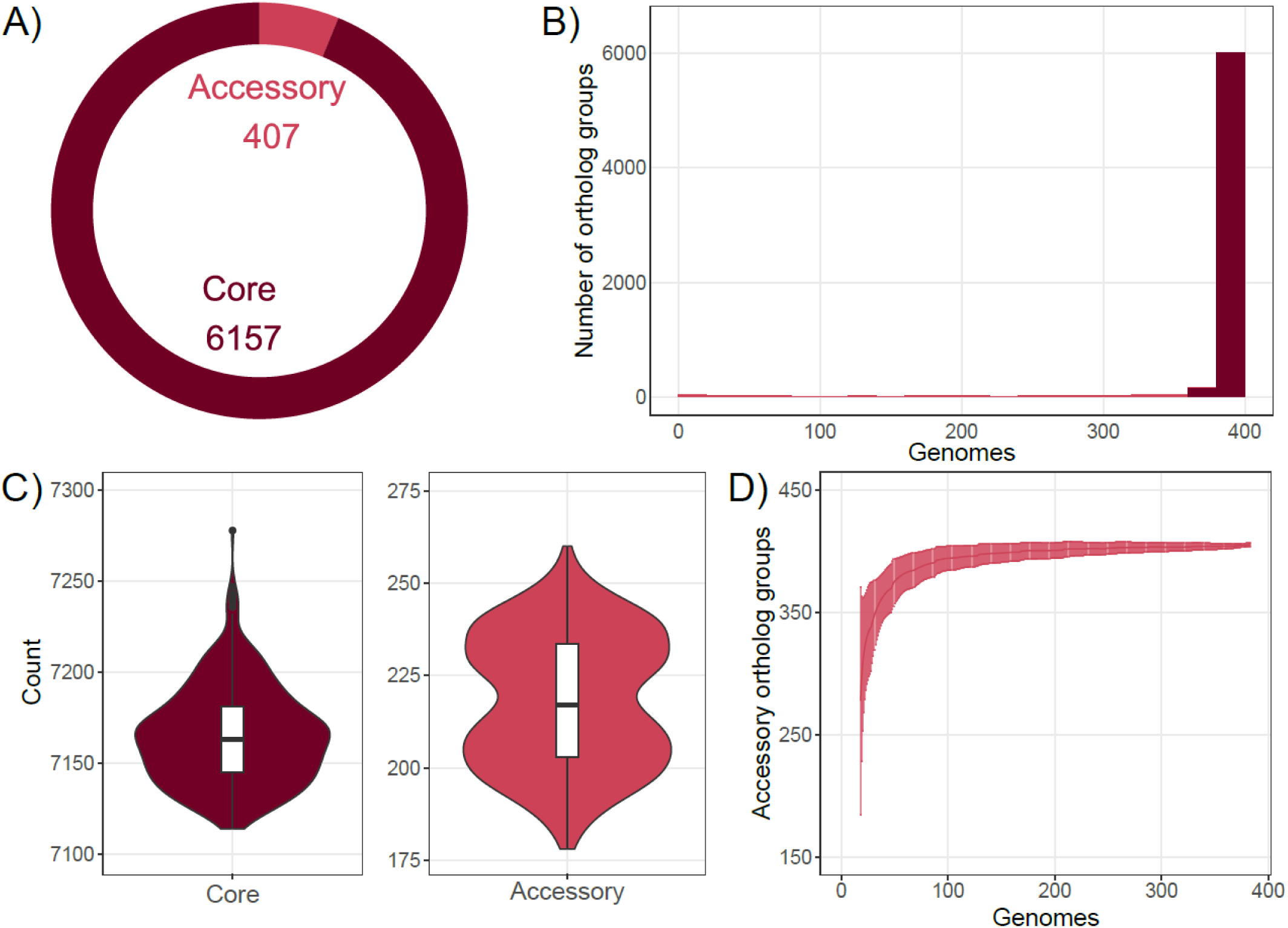
Distribution of core and accessory ortholog groups across isolates. **A)** Pie chart of 6564 ortholog groups assembled split into 2 classifications: core (6157 ortholog groups, 93.8% of total) and accessory (407 ortholog groups, 6.2% of total). **B)** Histogram of the number of ortholog groups and their presence total in unique genomes. Binned every 20 genomes. **C)** Violin plots with box and whisker plots of total core (left) and accessory (right) gene counts in each of the 383 genomes. **D)** Rolling mean and standard deviation every 20 genomes of 100 random samplings of the 383 genomes. Line indicates the number of accessory genes called as a new genome is added to the sample. Accessory ortholog groups were classified as <=95% of genomes currently in the sample (rounded).

### Cryptococcus neoformans pangenome is closed

Pangenomes can be classified as open or closed, based on how much of the true variation in gene content within a species is captured in the sample being examined. For example, adding an additional genome to an open pangenome has a high probability of adding novel genes. In contrast, in a closed pangenome additional genomes are unlikely to add novel genes. To determine whether the *C. neoformans* pangenome was open or closed, we sampled our pangenome one genome at a time and called the number of unique accessory ortholog groups found as a new genome was added. We did so by randomly sampling the 383 genomes 100 times (Figure 3d). Previously, bacterial pangenomes have been defined as open or closed based on a fitted equation to randomized sampling data using Heaps’ Law: *P=kN^γ^*where *P*=pangenome size, *N*=number of genomes, and γ is the new gene discovery rate[5]. Open pangenomes are those where additional sampling will add novel genes, while closed pangenomes are inferred to contain all the genes in the population. While γ>0 was originally used to classify a pangenome as open [5], more recently, a cutoff of γ<0.3 was proposed for a closed pangenome, with γ>0.3 indicating a pangenome that is still open[40,41]. For the rolling average of our accessory genes per genome, we fit the following equation: *P=*300.5*\*N*^0.05314^ (Figure 3d). With γ=0.05314, we can conclude that the *C. neoformans* pangenome is closed or nearly closed, depending on the cutoff chosen. We found nearly complete saturation, with 90% accessory ortholog groups (n=366) identified after the addition of only 42 random genomes to the pangenome.

### Molecular types harbor unique ortholog groups

The molecular type of *C. neoformans* isolates is defined using nucleotide similarity; therefore, we anticipated a high level of shared genomic content within each molecular type. We found that there were 14 VNI, 14 VNII, 4 VNBI, and 3 VNBII ortholog groups that were present only in their respective molecular types (Figure 4a). More frequently, however, we found ortholog groups that were present in three molecular types but entirely missing from the fourth, with 9 non-VNI, 97 non-VNII, 15 non-VNBI, and 6 non-VNBII ortholog groups (Figure 4a). Of note, ortholog groups not present in VNII are not always classified as accessory as we only included 15 VNII isolates in the population and our cutoffs for accessory content allowed them to be missing from 18 genomes and still be part of the core genome (n=10 non-VNII core ortholog groups) (Figure 4a). Overall, we found VNII to have the smallest number of shared ortholog groups when comparing molecular types to each other (Figure 4b). Some lineage-specific genes have previously been described, including the presence of *CNAG_06653* in VNI isolates[42]. We also found the orthologous cluster containing *CNAG_06653,* present in only 199/384 genomes, none of them from VNII isolates.

**Figure 4.**
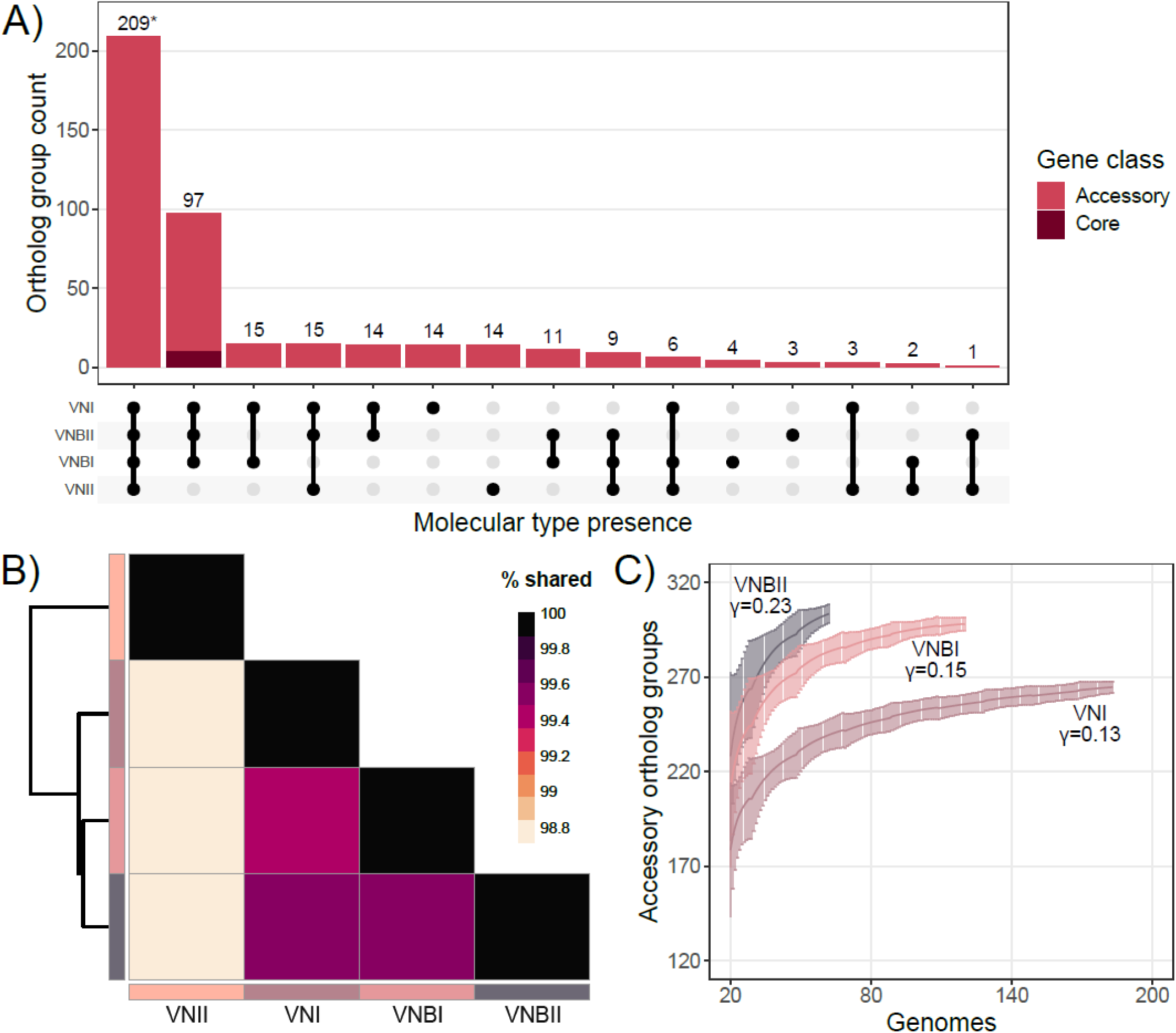
Molecular types offer unique accessory ortholog groups. **A)** UpSet plot of ortholog groups completely missing in at least one molecular type. Color indicates ortholog group classification as either accessory (pink) or burgundy (core). *First column shows all accessory genes that are not unique, i.e. distributed across all 4 molecular types. **B)** Analysis of shared ortholog groups between molecular types. Darker color (black) indicates a higher percentage of shared ortholog groups between compared molecular types. **C)** Rolling means and standard deviation of every 20 genomes of 100 random samplings of genomes belonging to a particular molecular type (n=184 VNI, n=121 VNBI, n=63 VNBII). VNII was not included due to only 15 genomes included in this collection.

To determine the gene diversity between molecular types, we also determined γ for each individually. We initially took 100 randomized samplings of genomes belonging to one individual molecular type and then calculated the rolling mean and standard deviation of new accessory ortholog groups for every 20 genomes across the 100 samples. Then, we fit a power equation to each molecular type to determine if pangenomes of each molecular type are open or closed. We found γ for each of the four molecular types: VNI=0.1343, VNBI=0.1528, and VNBII=0.228 (Figure 4c, Figure S4a, c-d). For VNII, since there were only 15 VNII isolates in this population, we calculated the rolling average and standard deviation every 5 genomes and predicted γ=0.403 (Figure S4b). From this information, we can infer that isolates from VNBII and VNII contain the highest probability of a new gene per new genome added to the population, with VNII considered open. However, the openness of a pangenome is heavily reliant on sample number, so increasing the number of VNII isolates may push the sample to being classified as closed. Further, as described above, the species pangenome is classified as closed, meaning that many of the “novel” genes that are added with additional sampling within a molecular type are genes already present in our pangenome from one of the other molecular types. Additional sampling thus may reduce the number of genes missing from a given molecular type but would not significantly expand the pangenome of the *C. neoformans* species as a whole.

### Essential genes are conserved

Because they are present in only some individuals within a species, accessory genes are less likely to have an essential biological role in standard growth conditions. To test whether core or essential genes were more likely to be absolutely required for growth, we took advantage of a resource previously developed by our lab. Briefly, our group classified genes within the reference strain H99 as essential or nonessential by determining their tolerance of random transposon insertions coupled with high-throughput sequencing (TN-seq)[43]. The vast majority of essential genes were annotated as core (n=1453, 99.6%), or present in greater than 365 of 384 total genomes. In contrast, only 0.4% of essential genes (n=5) were classified as accessory (Figure 5a). Accessory genes were far less likely to be classified as essential (*p*= 4.6e-09, Fisher’s exact test).

**Figure 5.**
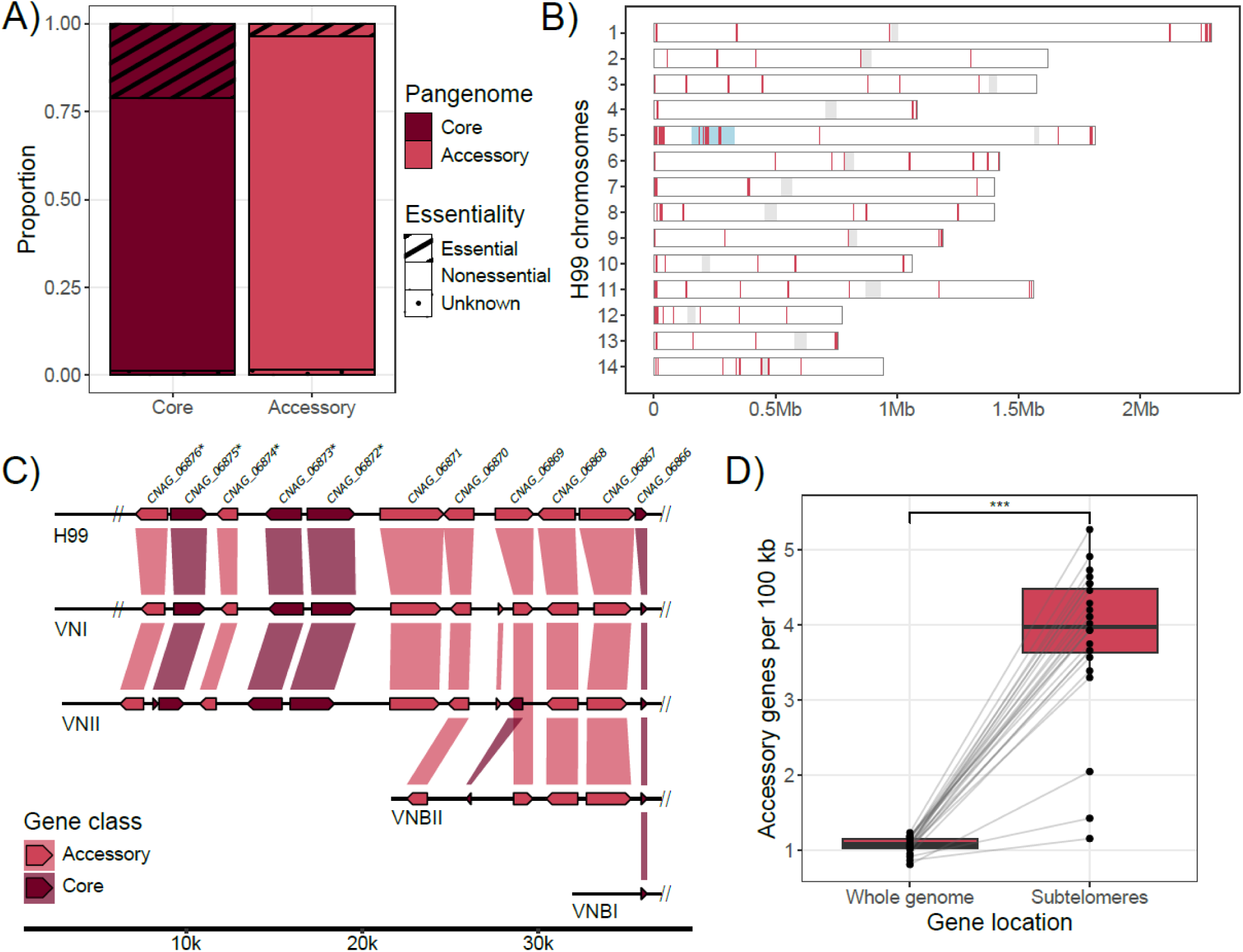
Accessory gene location across the genome. **A)** Stacked bar graph of proportion of core and accessory genes that were scored as essential, nonessential, or unknown in H99 TN-seq. Core genes were significantly more likely to be essential (*p*= 4.6e-09, Fisher’s exact test). 4 out of 5 total accessory essential genes were located within subtelomeres of H99. **B)** Map of 137 accessory genes along the 14 chromosomes of reference strain H99. Light grey circles indicate centromere position, light blue circles and pink lines indicate location of each accessory gene. Line width is representative of gene length. **C)** Closer look of accessory genes on the left arm of H99 chromosome 5. Each molecular type is represented along with the reference annotation from top to bottom with: H99 (reference, VNI), 125.91 (VNI), T4 (VNII), Bt1 (VNBII), and Ftc146-1 (VNBI). *CNAG* proteins are annotated on the H99 reference genome with *identity island genes. All sequences were aligned to edge of *CNAG_06866*. Hashes at the end of the genomes indicate further sequence, while no hash indicates end of contig. Base pair position on the bottom is relative to H99 genome chromosome 5 gene locations. Synteny bars between each gene indicates shared orthologous group while color indicates the ortholog group’s classification: core (burgundy) or accessory (pink). **D)** Box plot comparison of saturation of accessory genes located across the whole genome vs. subtelomeres in 24 long-read assemblies. Subtelomeres are defined as 40 kb of left and right arms of each chromosome assembled. Lines indicate each individual genome with points for the average number of accessory genes found per 100 kb. Paired *t*-test for subtelomeric > whole genome accessory genes per 10 kb (*t* = -14.454, df = 23, *p*-value = 2.481e-13).

Only five accessory genes were annotated as essential (*CNAG_00001*, *CNAG_05332*, *CNAG_06863*, *CNAG_07395*, and *CNAG_07654*). Four of these (all but *CNAG_06863*) were within 40 kb of the end of the chromosome, within a region often classified as the subtelomere[44]. *CNAG_06863* sits just outside of 40 kb from the end of the left arm of chromosome 5 and was deleted successfully in KN99[45]. Subtelomeric genes are known for high rates of recombination and gene loss, leading to potential differences between sequencing of the same isolate. A gene that is missing from an isolate used for a TN-seq experiment will be erroneously scored as essential because no transposons would map to that location in the reference genome. Additionally, subtelomeres are difficult to assemble with short read sequencing due to their repetitive nature across chromosomes. The location of these accessory essentials near the ends of chromosomes is reinforced by their slightly lower accuracy score (mean=0.89, n=5, all orthologs mean= 0.99).

### Accessory genes are located near telomeres

Previous analyses of the *S. cerevisiae* pangenome demonstrated that accessory genes were enriched within the subtelomeres[13], while other fungal pangenomes do not report similar enrichment[9,10,12]. To determine whether accessory genes are also concentrated within the subtelomeres of *C. neoformans*, we mapped all CNAG proteins that had been assigned to an ortholog group (n=6962) to the H99 reference available from NCBI (GCF_000149245.1). We visually saw that many accessory genes were located near the ends of the chromosomes. To test this, we counted the number of accessory genes within subtelomeres of H99. We again used the previous definition of subtelomeres for *C. neoformans*, 40 kb of the ends of the chromosomes[44], to evaluate accessory gene concentration. The defined subtelomeric regions (1.12 MB) comprised approximately 5.9% of the total genome of H99 (18.9 MB). We found that 65 of 137 (47.4%) accessory genes in the H99 genome were within our defined subtelomeres (Figure 5b).

Chromosome 5 of H99 harbored the highest number of accessory genes, containing 23 of the 137 total accessory genes (16.7%) (Figure 5b). Part of this can be attributed to the mating-type locus localized on the left arm of chromosome 5 (not subtelomeric). The *MAT* locus harbors several *MAT***a** and *MAT*α-specific genes that were annotated as accessory genes in the population. For example, the homeodomain transcription factors *SXI1*α and *SXI2***a** were classified into different accessory ortholog groups (CN06210 and CN06487). In addition, several *MAT***a** and *MAT*α alleles were sufficiently divergent to prevent mapping under our criteria. For example, *STE12* was annotated as an accessory gene despite it previously being annotated in both *MAT***a** and *MAT*α genomes[46]. Upon further investigation, *STE12* was not annotated by Funannotate in any of the 43 short-read *MAT***a** genome assemblies in our population and mapping representative *STE12* proteins from *MAT*α assemblies yielded no hits for gap-filling annotation with miniprot. Other accessory genes in the mating-type locus of H99 included *SXI1* (*MAT*α-specific gene), the *MAT*α pheromone, and smaller hypothetical proteins (*CNAG_06807*, *CNAG_06810*, *CNAG_01456*). Our short-read assemblies poorly assembled and annotated mating-type locus-specific genes, particularly in *MAT***a** genomes, contributing to the high number of accessory genes annotated on chromosome 5 of H99.

Mating-type loci aside, the subtelomeric region of the left arm of chromosome 5 also contains an “Identity Island” of genes with a high nucleotide identity (98.5%)[47] to the sister species *C. deneoformans* compared to the shared genome-wide nucleotide identity (88.17%)[33] (Figure 5c, Table 1). Hybrids between *C. neoformans* and *C. deneoformans* (AD hybrids) have no observable recombination along the identity island[48]. Within this region, we identified *CNAG_06876* as an accessory gene (Figure 5c). This accessory ortholog group was not present in any VNBI isolates (Table 1). *CNAG_06876* is influenced by calcineurin during temperature stress[49] and is annotated as an alpha-ketoglutarate-dependent taurine dioxygenase. Notably, all five genes within the identity island are either accessory or multicopy core ortholog groups.

**Table 1.** Identity island genes with high nucleotide identity between *C. deneoformans* JEC21 and *C. neoformans* H99. Ortholog group presence indicates average copy number of each gene and molecular types they are missing from.

| Annotation | JEC21 gene name | H99 gene name | Ortholog group presence in population |
| --- | --- | --- | --- |
| 5-oxoprolinase | *CNE05310* | *CNAG_06872* | Core (3-6 copies per genome) |
| 5-oxoprolinase | *CNE05320* | *CNAG_06873* | Core (3-6 copies per genome), grouped with *CNAG_06872* |
| HpcH/HpaI aldolase/citrate lyase | *CNE05330* | *CNAG_06874* | Accessory (0-1 copy per genome), not present in VNBI |
| Allantoate permease | CNE05340 | CNAG_06875 | Core (1-4 copies per genome) |
| Alpha-ketoglutarate-dependent taurine dioxygenase | CNE05350 | CNAG_06876 | Accessory (0-1 copy per genome), not present in VNBI |

To confirm that the relationship between subtelomeres and accessory genes was true outside of the reference strain H99, we examined accessory gene location for our non-reference long-read assemblies. Using only chromosome level assemblies (n=24), we mapped the locations of accessory genes for each isolate. Then, we determined if any portion of the accessory genes mapped fell within 40 kb of the ends of the assembled chromosomes. The subtelomeric regions were, on average, 5.9% of the assembled genomes. We found that per 100 kb, there was indeed a significantly higher density of accessory genes in the subtelomeric regions (Figure 5d, *p*= 2.481e^-13^, Fisher’s exact test). While slightly lower than the reference strain H99, we found that an average of 20.9% of accessory genes were located within the subtelomeres of non-reference isolates. The difference in proportion of accessory genes within subtelomeres between reference and non-reference isolates is most likely due to differences in annotation methods. Additionally, we annotated a higher number of genes, and thereby accessory genes, per genome at an average of 197 accessory genes mapped to long-read assemblies. Overall, there were still more accessory genes per 100 kb within subtelomeres (average=3.7 genes/100 kb) than the whole genome distribution (average=1.0 gene/100 kb) (Figure 5d).

### Accessory genes originate from related species

To determine where accessory genes originate from, we conducted a multispecies analysis within members of the Cryptococcaceae. We used miniprot to map all 407 accessory ortholog groups to 22 *Kwoniella and Cryptococcus* reference genomes we acquired from NCBI[46,50] (Figure 6a, Table S4). We required hits to cover at least 50% of the accessory query protein and reported the highest percent identity for each ortholog group in each species (Table S4). With these criteria, some orthologs did not map to any of the reference genomes outside of *C. neoformans* H99 (n=86). For example, ortholog group CN06219 contained reference protein *CNAG_07458* which has no predicted functional annotations and mapped to no other reference genomes from the Cryptococcaceae. The same was true for *CNAG_03972* and *CNAG_08014*. We note that this does not mean that accessory genes could not be segregating within each of those populations as well.

**Figure 6.**
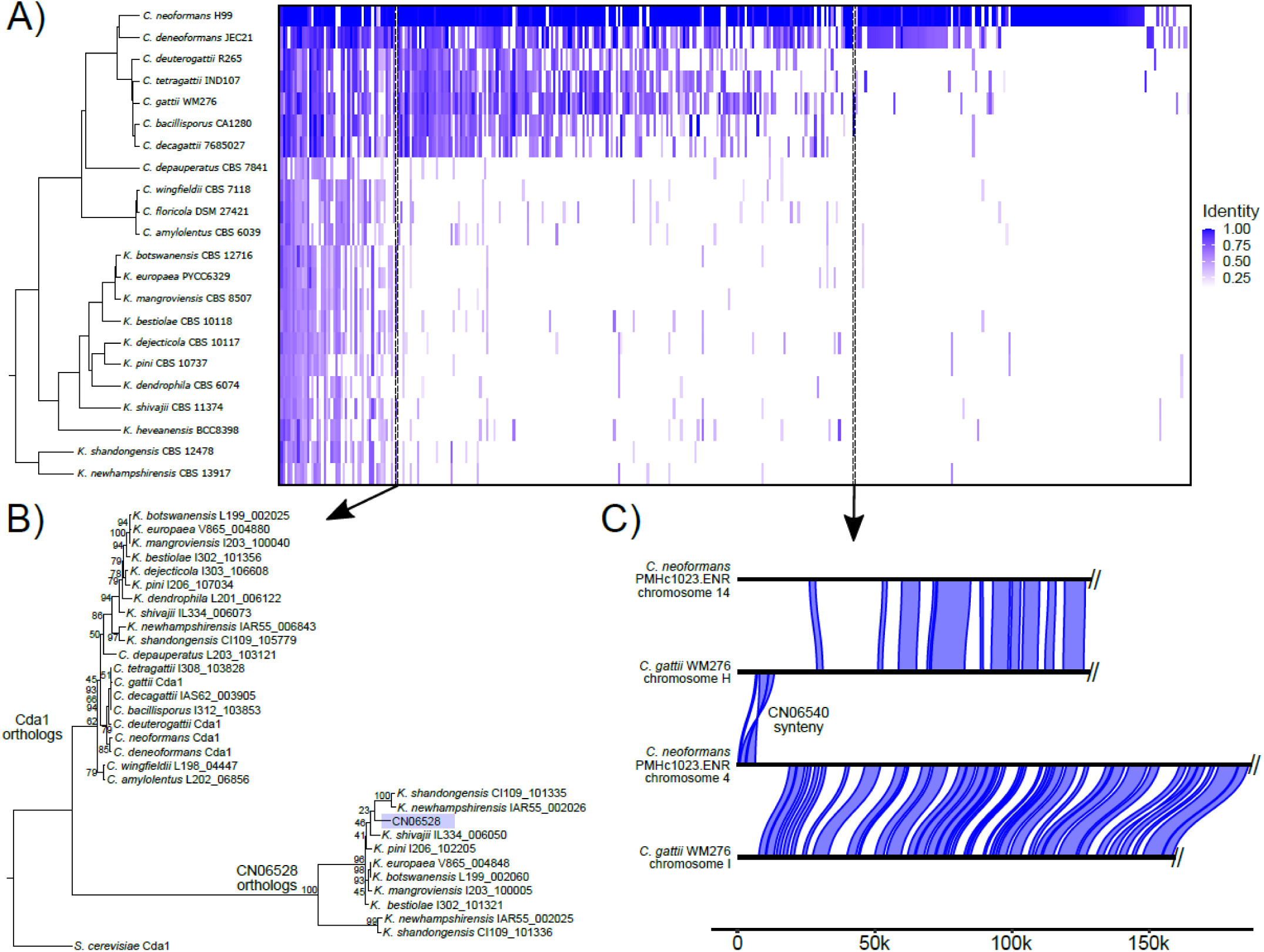
Accessory genes originate from related Cryptococcaceae species. **A)** Phylogenetic tree made by Colheo et al. 2025 (PLOS Biology) of type strains of *Cryptococcus* and *Kwoniella* species. Hits from miniprot mapping of accessory proteins (n=381) against assemblies of each species. Hits cover ≥ 50% of query accessory proteins and highest identity for each ortholog group species pair is represented by darker blue color. White indicates low identity or no hits; each column is an accessory ortholog group. Sorted by sum of identities across the species from highest (left) to lowest (right). Left dotted column correlates to ortholog group in panel B and right dotted ortholog group correlates to ortholog group in panel C. **B)** Tree of proteins containing similar accessory ortholog group CN06528 and Cda1 proteins. Proteins were aligned using MAFFT and 1,000 bootstraps and maximum parsimony tree made with RAxML. Percent bootstrap support shown on each node. Tree was rooted on model *Saccharomyces cerevisiae* Cda1. **C)** Synteny plot between ends of *C. gattii* WM276 chromosome H and VNII isolate *C. neoformans* PMHc1023.ENR chromosome 4 (not syntenic chromosomes). Above and below are true syntenic chromosomes for the pair (PMHc1023.ENR chromosome 14 & *C. gattii* chromosome H, PMHc1023.ENR chromosome 4 & *C. gattii* chromosome I).

The majority of the remaining 321 accessory ortholog groups mapped to the closest *C. neoformans* relative, *C. deneoformans* (n=232, 81% average identity) (Figure 6a, Table S4). Interestingly, the genome with the fewest mapped accessory ortholog groups was not the most distantly related, but instead was *C. depauperatus* (n=32, 51% average identity). *C. depauperatus* is a mycoparasite with a reduced genome size that may be the result of a shift to a parasitic lifestyle or loss of yeast phase growth[51]. These data suggest that many of the accessory genes segregating throughout the genus may have been lost during the genome contraction of *C. depauperatus*.

Interestingly, some accessory genes lacked identifiable orthologs in *C. deneoformans* but had orthologs in more distant *Cryptococcus* relatives. Many of these orthologs were present within the subtelomeres of these relatives. For example, ortholog group CN06397 was found variably in isolates of VNI, VNBI, and VNBII, but not of VNII. This ortholog group was highly similar to *CNBG_10092* from *C. deuterogattii*, a domain of unknown function containing protein located on the subtelomere of chromosome 11 in *C. deuterogattii*.

There are two potential rationales for a gene to have variable presence within a population. One is differential retention of genes, where a gene is present in the ancestral species but lost in some species over time (vertical gene transfer)[52]. The second rationale is horizontal gene transfer. Horizontal gene transfer events are evolutionarily more recent, where genes are directly transferred from one species to another through transposons, viruses, etc. We found both mechanisms are active in the *C. neoformans* pangenome. Below we describe an example of each using orthologs from the VNII molecular type.

CN06528 was found in all 14 VNII genomes and in 8/11 *Kwoniella* species. It contains a protein domain annotated as a chitin deacetylase (Cda1) but has been annotated in other fungal species as a polysaccharide deacetylase. Previously, *CDA1* has been identified as a potential antigen for cryptococcal vaccine development[53]. However, in H99 neither Cda1 nor any of its orthologs (Fpd1, Mp98), cluster with CN06528, indicating that this ortholog group is not a paralogous group resulting from a *C. neoformans* specific duplication of Cda1 (Figure 6b). In our long-read assembly of VNII isolate PMHc1023.ENR, we found the ortholog group within the right subtelomere of chromosome 8. Additionally, in the *Kwoniella bestiolae* genome, this chitin deacetylase resides on the left subtelomere of an assembled chromosome[32]. It is neighbored by another VNII-specific ortholog, CN06527, which also had orthologs only within 9/11 *Kwoniella* species, suggesting that both genes may have been segregating in a linked block. The average nucleotide identity between *K. shivajii* and *C. neoformans* PMHc1023.ENR genomes is 83.57% while the average nucleotide identity for CN06528 and the *K. shivajii* ortholog is 67.65%, suggesting there is not recent shared ancestry between these genes. These data are instead consistent with differential retention of the CN06528 ortholog group across the *Kwoniella* and *Cryptococcus* genera (vertical gene transfer).

CN06540 was present in only 8 VNII genomes but we identified highly similar orthologs in *C. gattii* and *C. bacillisporus*. In *C. gattii* WM276, *CGB_H0040W*[54] is located on the left subtelomere of chromosome H, which is not syntenic with reference *C. neoformans* strains. This ortholog family is predicted to encode a MviM dehydrogenase[55] containing a GFO/IDH/MocA domain[32] but is otherwise of unknown function. In our long-read assembly of VNII isolate PMHc1023.ENR, we found the member of this ortholog group within the subtelomere of chromosome 4. Upon mapping of these two chromosomes, which are not syntenic, we found 5,768 bp of shared synteny between the two subtelomeres (Figure 6c). The average nucleotide identity for CN06540 and the orthologous *C. gattii* gene *CGB_H0040W*, however, was 97.75% compared to the genome-wide average nucleotide identity of 86.04%. The high identity between genes suggests horizontal gene transfer between *C. gattii* and VNII genomes. The syntenic region encompassed three genes in PMHc1023.ENR: ortholog group CN06540, CN00037 (core myo-inositol 2-dehydrogenase), and CN03105 (core BNR/Asp-box repeat family protein). In both genomes, the region was immediately next to or contained a long terminal repeat region found in *Cryptococcus* (LTR13), suggesting that either this homology or movement of a retrotransposon may have facilitated this genetic transfer.

## DISCUSSION

Here we describe the first large scale pangenomic analysis of *C. neoformans* isolates, to our knowledge. We *de novo* assembled and annotated 383 diverse *C. neoformans* isolates[17] spanning all 4 molecular types to generate a pangenome of 6,564 orthologous groups for the population. We generated high-quality assemblies and annotations from Illumina short-read sequencing that we benchmarked by long-read sequencing for 9% of isolates. This is in contrast to earlier efforts in *C. neoformans* that used smaller numbers of genomes and lacked methods to benchmark accessory gene accuracy. The first *C. neoformans* pangenome was based on 25 *C. neoformans* genomes[6]. It contained 5,486 core genes and 964–1590 accessory genes, with a total of 80.56% of orthologs as core[6]. In comparison, our pangenome identified a larger number of orthologous groups (1,078 more) but annotated fewer as accessory (407 accessory ortholog groups). We found the *C. neoformans* population we used for pangenome construction was nearly closed with a low rate of novel genes found per additional genome.

We determined that each of the four molecular types in *C. neoformans* had both shared gene and unique genomic content. The diversity between molecular types was exemplified by the gene content we found on the left arm of chromosome 5, where subtelomeric segments appeared to vary in length with corresponding differences in accessory gene content. The genomic content differences we found between molecular types aligns with previous work, showing that despite their shared species, each molecular type harbors specific SNPs[17]. We found VNII to have the highest rate of new genes found per genome added and to have the lowest number of shared ortholog groups with the other molecular types. We found that most accessory ortholog groups were shared between VNI, VNBI, and VNBII, leaving 97 non-VNII ortholog groups. Additional sampling of VNII will likely improve future iterations of the *C. neoformans* pangenome. While VNII-specific accessory genes were the largest group of genes unique to a molecular type, we found that all four molecular types harbored their own unique ortholog groups. These data are consistent with previous observations of limited recombination and thus restricted gene flow between the *C. neoformans* molecular types[17].

We found that *C. neoformans* has similar content diversity to other budding yeasts. Pangenomes have been generated for both pathogenic (*Candida albicans*[6]*, C. parapsilosis*[10]*, C. auris*[12]*, Nakaseomyces glabratus*[56]) and nonpathogenic species (*S. cerevisiae*[13]) of budding yeasts. In general, budding yeasts have a smaller proportion of accessory ortholog groups (63.3[13] –99.2%[10] core) than filamentous fungi, like *A. fumigatus* (55.42[9]–69%[11] core) and *Zymoseptoria tritici* (58.1% core[57]). Filamentous fungi have fundamentally different lifestyles than yeasts, including behaviors like vegetative hyphal fusion that can place nuclei from genetically distinct isolates into a shared cytoplasm. As a result, they may be more prone to horizontal gene transfer between isolates, increasing the likelihood that newly developed or acquired accessory genes spread through a population. Both *A. fumigatus* and *Z. tritici* are also host to *Starship* transposable elements[15,58], which could facilitate movement or introduction of novel accessory genes, while this class of giant transposable element is not found in any of the yeasts described above.

Fungal genomes vary not just in the amount of the genome that is accessory, but also in the locations of those accessory genes. In both *C. auris*[12] and *A. fumigatus*[13], accessory genes are distributed throughout the genome rather than in particular enriched regions. For *A. fumigatus* it is thought that large *Starship* transposons are moving most of the accessory genes to various locations along the genome[13], but native *C. auris Starships* have not been identified. In contrast, we found that accessory genes were enriched within the subtelomeres, which is similar to previous observations in the nonpathogen *S. cerevisiae*[13]. Accessory genes and structural variation in the subtelomeres of *S. cerevisiae* were driven by horizontal gene transfer events from divergent yeast species[13]. Likewise, we found many accessory genes in the *C. neoformans* subtelomeres were from related Cryptococcaceae species. Therefore, the concentration of subtelomeric accessory genes may be driven by horizontal gene transfer events with other fungi. The origin of horizontal gene transfer events in *S. cerevisiae* is thought to be from fungal species that reside in similar fermentative environments[13]. Similar logic would suggest that Cryptococcaceae species residing in similar environments together, like trees[59], may be able to exchange subtelomeric genes across species barriers via horizontal gene transfer rather than sexual reproduction. These data also suggest that *S. cerevisiae* and *C. neoformans* may share horizontal gene transfer mechanisms that enable subtelomeric gene acquisition that are missing from *C. auris* and *A. fumigatus*. Alternately, perhaps genome defense mechanisms prevent incorporation of foreign DNA into chromosome bodies more effectively than into the subtelomeres in *C. neoformans* and *S. cerevisiae* but not in *C. auris* and *A. fumigatus*. Future experiments will be necessary to explore these hypotheses.

There are two *C. neoformans* isolates (Bt65 and Bt81) that lack RNA interference (RNAi) machinery, which results in increased insertions of the Cnl1 retrotransposon near telomeric repeats[39]. RNAi is a genomic defense mechanism that helps prevent transposon movement in *C. neoformans*, thereby limiting the copy number of transposons in the genome[60]. Interestingly, we found these two RNAi-defective genomes to have the two lowest number of accessory genes mapped to the subtelomeres of the long-read sequences included in this study (Bt65: 16 accessory genes, Bt81: 13 accessory genes, average: 42.75 accessory genes in subtelomeres). Since we define subtelomeres as the last 40 kb of each chromosome and because Cnl1 elements comprise much, if not all, of this region in Bt65 and Bt81, the relative lack of accessory gene enrichment is primarily driven by lack of gene annotation within the subtelomeres. Therefore, we conclude that unrestricted transposon activity is unlikely to be a strong positive contributor to content variation in the subtelomeres of *C. neoformans* and, if anything, acts to limit accessory gene accumulation in this region.

Subtelomeres are important for environmental adaptation as well as pathogenic immune evasion. In the human fungal pathogen *N. glabratus*, the subtelomeres harbor adhesion-related genes which allow the pathogen to change its cell surface and evade host immunity[61]. Similarly, the fungal pathogen *Pneumocystis carinii* subtelomeres contain major surface glycoproteins[62] which are used to evade immune responses through differential expression[63]. Parasitic pathogens like *Plasmodium spp.* and *Trypanosoma brucei* also contain diverse repertoires of 600[64]-2,000[65] surface antigen glycoproteins within subtelomeres. These parasites can evade host immunity by altering which surface antigen is expressed. *C. neoformans* contains numerous annotated sugar transporter genes within its subtelomeres[44,66], potentially contributing to cell surface changes as seen in other eukaryotic pathogens. It remains to be seen whether subtelomeric accessory genes in *C. neoformans* contribute to growth in the environment, virulence, or both. Future experiments exploring the phenotypic consequences of gene content variation and subtelomeric accessory gene variation, in particular, will be necessary to elucidate the impact of these genes on infection and disease progression.

## MATERIALS AND METHODS

Isolates used in this study were a part of the Strain Diversity Collection assembled by Desjardins et al. under NCBI BioProject PRJNA382844 (Table S1) [17]. These isolates encompassed both *MAT***a** and *MAT*α mating types across the four identified molecular types for *Cryptococcus neoformans* var. *grubii*, since renamed *C. neoformans*[67]. Isolation sources include both clinical and environmental sources, creating one of the most diverse collections of *C. neoformans* isolates.

### Short-read assembly

We downloaded all Illumina sequence runs for 387 samples from Desjardins et al.[17] using NCBI’s SRA-Toolkit v3.0.1[50] (2025-06-09). This resulted in a total of 1701 downloaded runs with an average of 4.39 runs per sample. Next, we used AAFTF v0.5.0[68] to trim, filter (BBMap v38.84[69]), filter, and assemble (SPAdes v3.13.1[70]) each genome from our downloaded reads. We installed AAFTF as a Bioconda v24.11.3[71] environment. We removed duplicate contigs (minimap2 v2.22[72]) and polished (Pilon v1.24[73]) assemblies. Finally, we scaffolded our assemblies to the H99 reference genome using RagTag v2.1.0[74] to make longer contigs from assemblies. We kept contigs that did not align to the reference genome for subsequent analysis. Our finalized assemblies were assessed using AAFTF[68] *assess*. We excluded any assemblies with >200 contigs from further analysis (n=3: Bt206, Bt92, and PLR10).

### Short-read annotation

Our annotation of our assemblies was carried out using Funannotate v1.8.15[75]. First, we sorted our assemblies by length using Funannotate *sort*. Next, we masked repeat regions of the assemblies using TANTAN v51[76].We supplied RNA-seq data from NCBI’s BioProject PRJNA167480[20] from reference strain H99 to guide Funannotate. These included two replicate RNA-seq runs of H99 grown in three separate conditions: YPD, pigeon guano broth, and starvation media[20](Table S1). We then used Funannotate *train* with flags “*--jaccard_clip --pasa_db mysql*” to run Trinity v2.8.5[77] to assemble the transcripts from the 6 RNA-seq runs (n=16,765) to provide as evidence for gene prediction. PASA v2.5.3[78] then generated gene models from the RNA-seq data which were fed into Kallisto v0.46.1[79] to select a model for each locus.

We then used transcript evidence to predict genes in the assemblies using Funannotate *predict* with flags *“--min_protlen 35 --repeats2evm --keep_nostops --augustus_species cryptococcus* “. Additionally, we provided Funannotate with H99 annotated proteins and transcripts downloaded from FungiDB v68[32] “FungiDB-68-CneoformansH99” (2025-06-09). Funannotate *predict* used 5 *ab-initio* predictors to supplement transcript evidence: AUGUSTUS[80] v3.5.0 (pretrained with *Cryptococcus* data), CodingQuarry v2.0[81], GeneMark-ET v4.71[82], GlimmerHMM v3.0.4[83], and SNAP v2017_03_01[84].

We annotated our final gene set for predicted function using Funannotate *iprscan* implementing InterProScan v5.63-95.0[85]. Additionally, we predicted transmembrane regions of proteins using Phobius v1.01[86]. Funannotate *annotate* also used UniProt DB v2023_05[87], Pfam v36.0[88], and eggNOG-mapper v2.1.12[89,90] to annotate parsed proteins. We used the BUSCO v2.0[91] basidiomycota_odb9 (2016-02-13) database to assess the quality of our annotations. We excluded genomes that had <96% of the 1,335 BUSCOs proteins searched (n=1: WM626.MMRL.3150).

### Ortholog group construction

To make orthologous clusters, we fed our protein sequences to OrthoFinder v2.5.5[92]. We also included the longest protein sequences (n=6,976) of the longest single transcripts of reference strain H99 accessed from FungiDB v68[32]. We used the “*-op*” command for OrthoFinder to run DIAMOND v2.1.8[93] separately for protein-protein comparisons. Then, we finished the OrthoFinder analysis by using *“-b”* command.

### Ortholog group validation

After initial construction with OrthoFinder[92], we went through several steps to validate orthologous clusters (n=9,194) of protein sequences. First, we checked ortholog groups for accidental combination of genes by either missing stop codons or annotation errors using CNAG protein annotations for alignment. This resulted in the separation of 15 ortholog groups (new ortholog group count: n=9,186). Then, we manually removed all contaminants (n=37) and singletons with no start codon (n=24). Next, we used BLAST+ v2.13.0[94] to identify ortholog groups that were highly similar to each other by aligning the longest representative protein of ortholog groups present in >20 unique genomes against ortholog groups present in <=20 genomes. Ortholog groups found in <=20 genomes with >80% identity to a larger ortholog group were combined with that larger ortholog group (n=1,027).

To further verify our ortholog groups, we manually investigated all ortholog groups against reference strain H99 proteins. We used IGV v2.18.2[95] to view BLAST alignments of the longest ortholog group representative sequence against H99 protein sequences with e-values < 1e-05. We manually investigated all ortholog groups containing ≤ 365 unique genomes. We then combined ortholog groups that were highly similar to H99 proteins with the representative clustered H99 protein ortholog group (n=459).

To ensure we were not missing any unannotated proteins, we used miniprot v0.13[96] to annotate genomes using existing orthologous clusters. We selected representative proteins of current missing (n<383 unique genomes) orthologous clusters by selecting the longest H99 reference protein. If there is no reference protein, we selected the longest protein in the cluster. Then, using the set of representative proteins, we used miniprot to align proteins to previous assemblies for each isolate. We selected alignments that had >75% identity and >80% of the representative protein aligned. We then used gff files generated by Funannotate to identify if newly mapped proteins were overlapping with established annotations. This resulted in 631 unique pairs of overlapping ortholog groups in the 383 assembled genomes. We combined any two ortholog groups that had overlapping map sites in >90% of total genomes in the orthologous cluster (n=133). Using our final ortholog group set, we again used miniprot to annotate genomes for proteins that were: 1) not present in the genome, 2) had 75% identity to the representative ortholog group protein sequence, 3) aligned to >80% of representative ortholog group protein sequence, and 4) did not overlap with previously annotated genes via Funannotate (n= 172,423 proteins, n= 1,162 ortholog groups).

To combine all reference protein ortholog groups, we reclustered ortholog groups (n=792) using FungiDB[32] OrthoMCL v6.21[97] (2026-03-04) ortholog clusters (n=6,067). We then manually checked if accessory reference proteins were correct or if there were annotation errors in the remaining 383 genomes. This led to manually splitting 12 more ortholog clusters using H99 reference proteins to separate combined annotated proteins. Additionally, we manually added fragmented proteins to ortholog groups that were known to be conserved but were not called due to scaffolding (n=4). Unknown amino acids in these proteins were labeled with an “X”. All ortholog groups containing reference proteins belonging to the mitochondrial genome were manually removed (n=12). We finalized our ortholog group count at 6,564 ortholog groups, which were then clustered into core (>95% of genomes) or accessory (≤95% of genomes)[9]. The core cutoff for our 384 genomes was >365 genomes (Table S2).

### Long-read DNA extraction, sequencing, and assembly

We used a DNA extraction protocol that was modified from Pitkin et al.[98,99]. We grew isolates (n=10) from single colonies in 100 mL of yeast peptone dextrose (2% Bacto Peptone, 1% Bacto Yeast Extract, and 2% glucose) media shaking for 24 hours at 220 rpm. We then lyophilized our cultures overnight and used 3-5mL of 3mm glass beads to break apart cells by vortexing. Then, we suspended cells in 20 mL of extraction buffer (100 mM Tris-HCl, 0.7 M NaCl, 0.5 M EDTA, 1% cetyltrimethylammonium, 1% beta-mercaptoethanol). We then incubated the cells at 65°C for 1 hour and then cooled on ice for 10 minutes. After chilling, we added an equal volume of chloroform and pelleted via centrifugation at 3900 rpm for 10 minutes. We moved the supernatant into a fresh 50 mL conical tube where we then added an equal volume of isopropanol and incubated at -20°C for 1 hour. We pelleted the DNA via centrifugation at 3,900 rpm for 10 minutes at 4°C and washed with 70% EtOH. We then suspended the DNA pellet in 1 mL TE (10mM Tris and 1mM EDTA, pH 8) with 100 µg/mL RNAse and incubated at 37 °C for 30 minutes. We then added a final volume of 0.5 M sodium acetate solution to precipitate the DNA. We went through a second round of adding equal chloroform, centrifuging at 13,000 rpm to pellet, removing the supernatant to a clean tube, and adding an equal volume of isopropanol. Finally, we spooled the DNA into a clean tube, washed with 70% EtOH, and suspended in 200 µL of 1X TE.

We then sequenced our long-stranded DNA through a commercial provider (Plasmidsaurus) using Oxford Nanopore Technologies (ONT) on a PromithION with R10.4.1 flow cells using Dorado dna_r10.4.1_e8.2_400bps_sup@v4.3.0 to call bases. We then used Canu v2.3[100] with “*contigFilter=”2 200000 1.0 0.5 5””* to filter out contigs < 200 kb and “*genomeSize=19m correctedErrorRate=0.075”* to assemble our ONT sequences into larger contigs (Table S3). After Canu assembly, we polished with long-read sequences using Medaka v1.12.1 (https://github.com/nanoporetech/medaka) with *medaka_consensus*. We polished our assemblies again with available Illumina sequences for each genome under PRJNA382844. We first used minimap2 v2.24[72] to align short-read sequences to our draft assembly, then used Pilon v1.24[73] to iteratively polish our draft assemblies 5 times, aligning to the new assembly after each round of polishing[101].

We assessed the quality of long-read assemblies by mapping back to the H99 reference genome (from FungiDB[32]) with minimap2 and visualized alignments with pafr (https://github.com/dwinter/pafr). We renamed and oriented contigs to align with H99 chromosome names. We named contigs that had alignments to several chromosomes based on which contig had the most coverage on a given chromosome or which contig contained the H99 centromere if indistinguishable. Additionally, we scaffolded any contigs that had broken at the rRNA locus on chromosome 2 with 50 Ns[101]. We looked for telomere repeats[102], 5′-TAACCCCC-3′, at the ends of our contigs. We confirmed each isolate was *C. neoformans* by aligning the H99 *URA5* locus to each assembly and filtered those with poor sequence similarity (n=1). Long-read sequences and assemblies are available under PRJNA1477025 (Table S3).

We downloaded additional publicly available long-read assemblies (n=26) [33–39] of isolates within our collection of 383 genomes using NCBI Datasets command-line tools v18.26.0[55] (Table S3). All long-read assemblies were assessed for completeness using BUSCO v6.0.0[91] and the basidiomycota_odb12.2 dataset (Table S3). We filtered out any assemblies with a complete BUSCO score < 96% (n=1, C8).

### Long-read annotation and ortholog group validation

We used our 35 long-read assemblies to validate if missing ortholog groups were due to gaps in assembly or true gene loss. To do so, we used protein annotations from both Funannotate and miniprot to align to long-read assemblies. All annotations that were generated from Funannotate and miniprot annotation of short-read assemblies were mapped to long-read assemblies. Any ortholog groups that were not called via short-read assembly annotation were added for mapping with a representative sequence that was either the longest CNAG protein or longest protein. Similar to our initial miniprot annotation, we only called ortholog groups as a hit if they covered 80% of the query ortholog group and had >75% identity.

We then determined the accuracy of our short-read ortholog group annotations using a confusion matrix (Figure S2a). Ortholog groups that were mapped in both short and long-read assemblies were true positives and ortholog groups that were not mapped in either were true negatives. Ortholog groups that were annotated in short-read assemblies but not in long-read assemblies were false positives. Finally, ortholog groups that were mapped in long-read assemblies but not in short-read assemblies were false negatives. Copy number was not considered in the accuracy for individual ortholog groups, only presence or absence of a given ortholog group in the 35 genomes.

### Ortholog group analysis and visualization

Figures were made with ggplot2 v4.0.3[103] on R v4.6.0[104]. Final figures were all assembled using Inkscape v1.1.1. We used the R package pheatmap v1.0.13[105] to create heatmaps of shared accessory genes and ggupset v0.4.1[106] to create an upset plot. The R package gggenomes v1.1.3[107] was used to visualize chromosome 5 subtelomeres across molecular types and synteny between species. To examine species accessory hits, we used package ggtreeExtra v1.22.0[108] to generate a heatmap.

We used MAFFT v7.526[109] to create alignments for CN06528 from amino acid sequences acquired from NCBI[55] and FungiDB[32] on 2026-09-22. Using CN06528 alignments, we then inferred a maximum likelihood tree using RAxML v8.2.13[110] using 100 inferences of randomized maximum parsimony trees with “*raxmlHPC -m PROTCATWAG -#100*”. We then conducted 1,000 bootstraps using “*raxmlHPC -f d -# 1000*”. Bootstrap percentages were added to the best-fit tree using “*raxmlHPC -f b”*. We then visually examined the tree using MEGA v11.0.13[111] and then used ggtree[112] along with ggtreeExtra v1.22.0[108] to create the final phylogeny.

To compare synteny across alignments we used minimap2 v2.29[72] with flags “*-cx*”. For comparisons between isolates, we also used argument “*asm5*” but for between species we used “*asm20*”. To calculate average nucleotide identity between isolates we used pyani v0.2.11[113] “*average_nucleotide_identity.py*”. To compare average nucleotide identity between genes, we used the flag “*-m ANIb”*.

## DATA AVAILABILITY

Code used is available at: https://github.com/Claire-Reichardt/C.neoformans_Pangenome.git. Long-read sequencing and assemblies generated in this study are available under BioProject PRJNA1477025. Short read assemblies and annotations are available at 10.5281/zenodo.23697234.

## SUPPLEMENTARY DATA

1. **S1 Table. Illumina sequence assembly and annotation information. A)** List of 387 isolates used as reported by Desjardins et al. 2017[17]. **B)** RNA-seq evidence from H99 used to annotate short read assemblies from Janbon et al. 2014[20]. **C)** Annotation and assembly statistics for each isolate generated by Funnannotate. **D)** BUSCO statistics for each isolate. **E)** Final count of core and accessory genes called for each isolate.
2. **S2 Table. Ortholog group information. A)** List of final ortholog groups generated along with classification, count, copy numbers, and accuracy scores. **B)** CNAG proteins and their classified ortholog group. **C)** Longest non-CNAG protein representative sequence for ortholog groups. **D)** Number of proteins in each ortholog group for each isolate.
3. **S3 Table. Long-read assembly and annotation information. A)** 34 long-read assemblies accessed from NCBI and generated in this study [33–39]. **B)** BUSCO scores for long-read assemblies. **C)** Mapping accuracy for short-read ortholog groups to long-read assemblies. **D)** Accessory genes mapped to subtelomeres of long-read assemblies.
4. **S4 Table. Species hits to accessory orthologs. A)** Table of all downloaded Cryptococcaceae species representatives from NCBI. **B)** Highest reported miniprot identity from accessory protein to species genome alignment that covers ≥50% of query protein.

## ACKNOWLEDGMENTS

We thank members of the Billmyre lab and Katie Billmyre for helpful comments on the manuscript. We would also like to thank Teresa O’Meara and Zoë Hilbert for their helpful conversations in the development of this manuscript.

## SUPPLEMENTARY FIGURES

**Figure S1.**
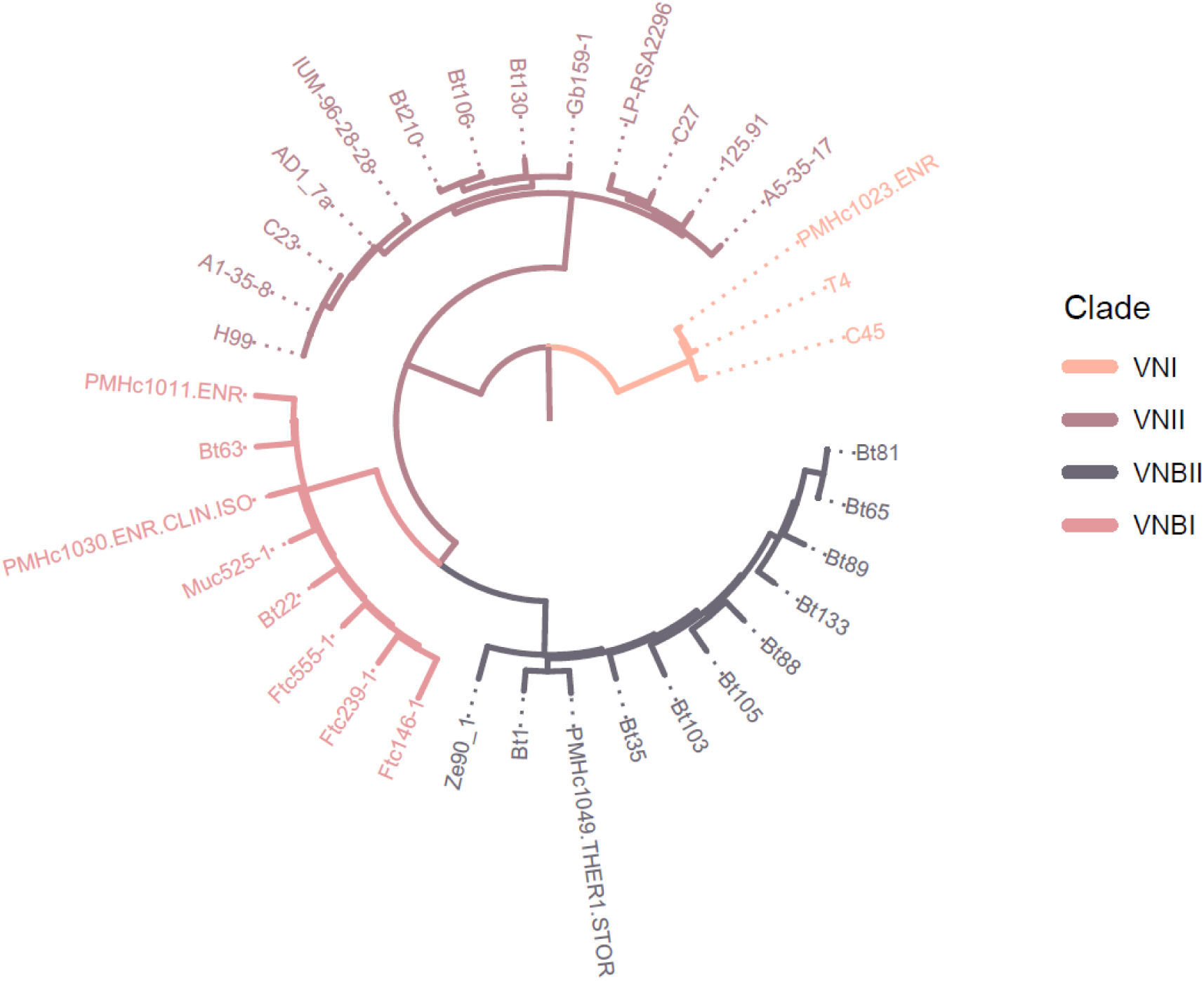
Phylogenetic tree of long-read assemblies included in this study. Tree was calculated by Desjardins et al. 2017 and rooted on VNII.

**Figure S2.**
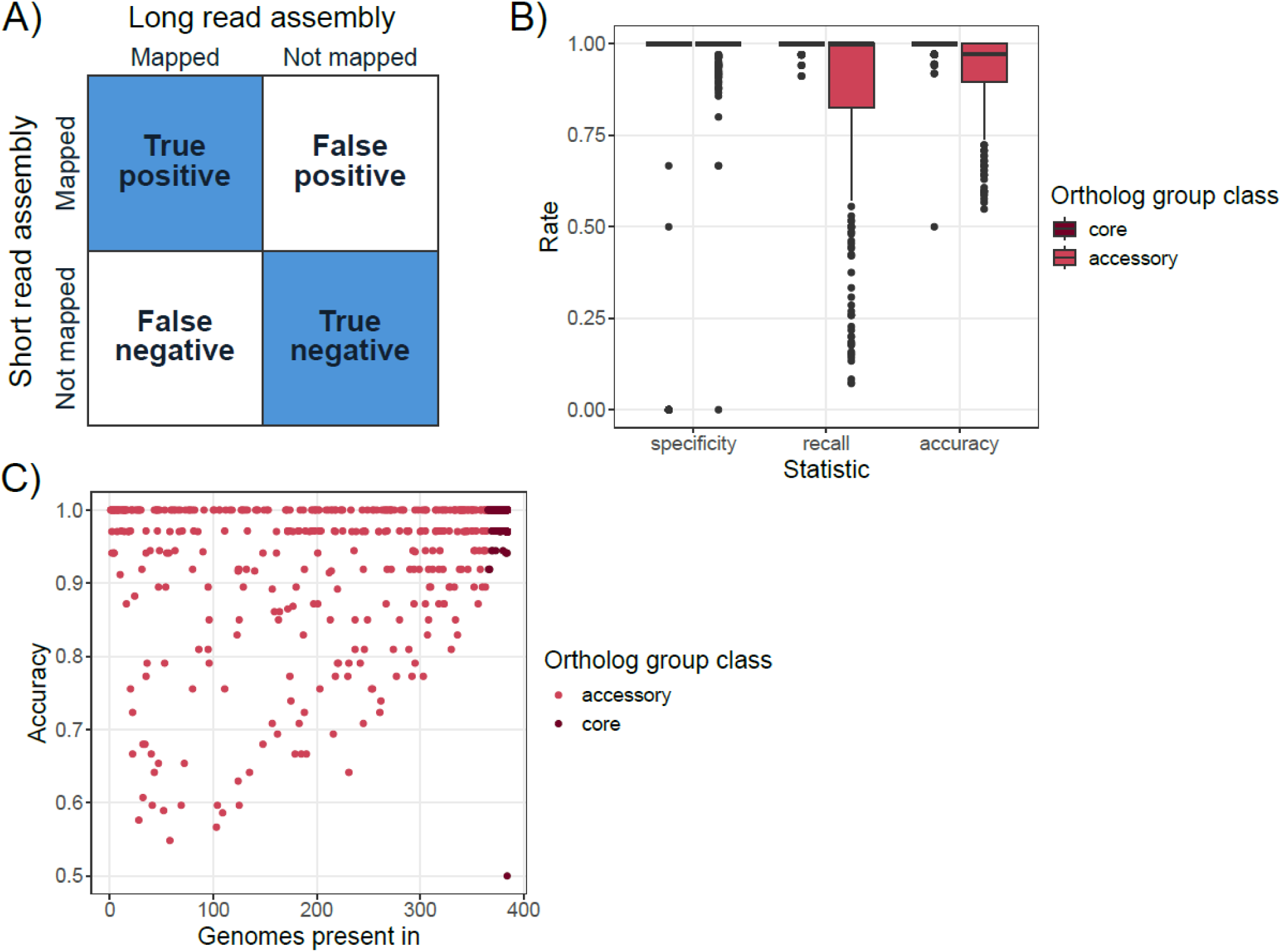
High accuracy of orthologs mapping to long-read assemblies. **A)** Confusion matrix determining accuracy of short-read assembly ortholog presence/absence compared to long-read assembly mapping data. **B)** Box and whisker plot of summary statistics of core and accessory ortholog groups from mapping short-read annotations to long-read assemblies. Plot shows specificity (1-false positive rate), recall (1-false negative rate), and accuracy for core and accessory ortholog groups. **C)** Ortholog group accuracy compared to the number of genomes that have a given ortholog group.

**Figure S3.**
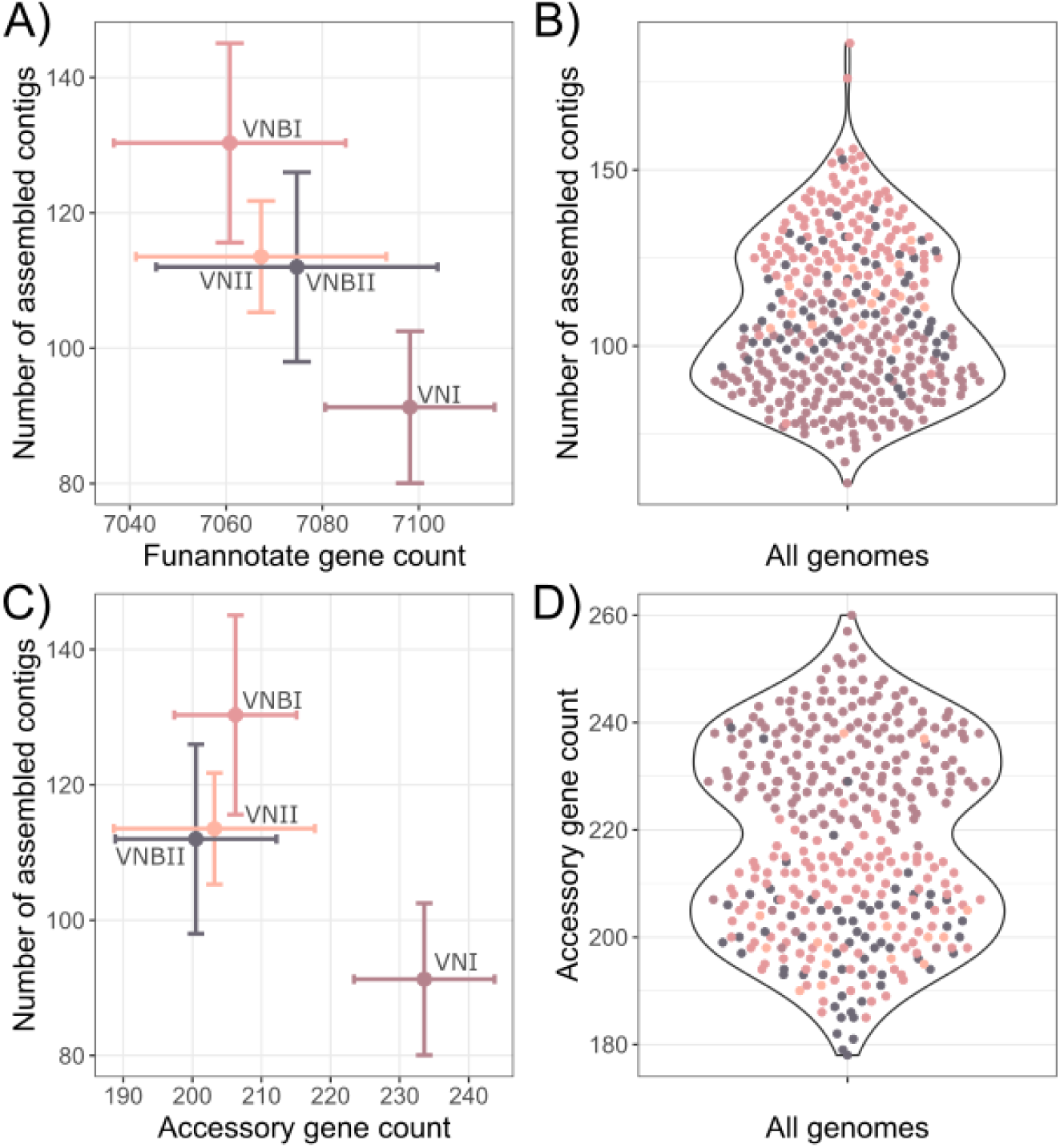
Accessory gene count correlates with assembly quality. **A)** Direct number of genes called for each molecular type after Funannotate annotation before manual curation. Points show the average for each group and bars are the standard deviation. **B)** Number of assembled contigs per genome. **C)** Final accessory gene count vs. the number of assembled contigs per each molecular type. **D)** Final accessory gene count with points annotated per genome colored by molecular type assignment.

**Figure S4.**
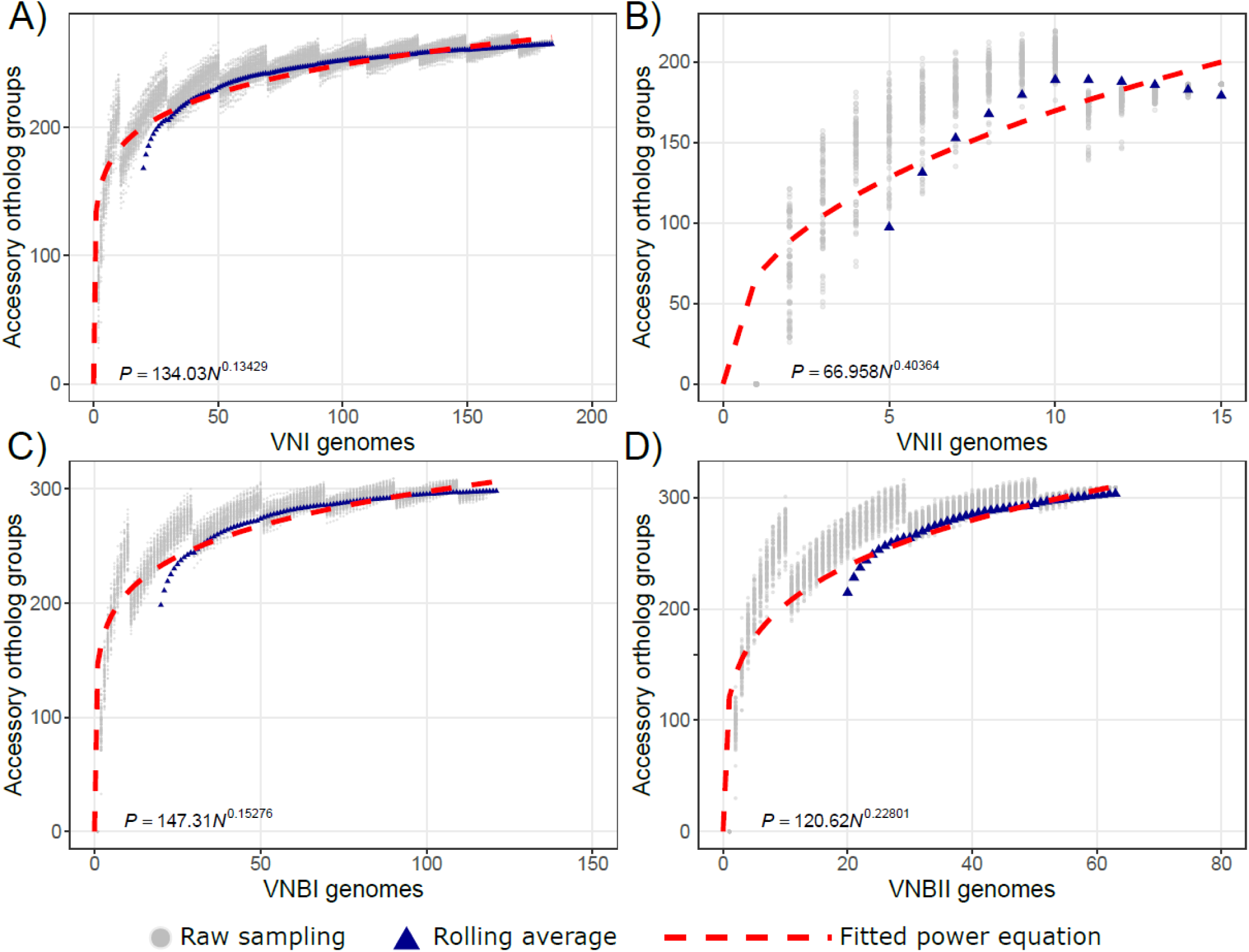
Determining the discovery rate of new genes for each molecular type. 100 randomized samples for the number of new accessory ortholog groups found for each new genome added (grey dots) were taken for each genome within a particular molecular type. Sample sizes were: 184 VNI, 15 VNII, 121 VNBI, and 63 VNBII isolates. Power equations (bottom of graph) were fit (red doted lines) to the rolling average for a given molecular type (blue triangles). **A, C-D)** rolling average was calculated every 20 genomes while **B)** was calculated every 5.

